# Characterizing noncompliance in conservation: a multidimensional Randomized Response Technique for multinomial responses

**DOI:** 10.1101/453159

**Authors:** Jacopo Cerri, Lapo Scuffi, Annamaria Nocita, Marco Zaccaroni, Andrea Lenuzza, Maarten Cruyff

**Affiliations:** Istituto di Management, Scuola Superiore Sant’Anna. Piazza Martiri della Libertà 33, 56127, Pisa, Italy; Istituto Superiore per la Protezione e la Ricerca Ambientale (ISPRA). Piazzale dei Marmi 2, 57123, Livorno, Italy; Scuola di Scienze Matematiche, Fisiche e Naturali, Università degli Studi di Firenze. Viale Morgagni 40/44, 50134, Firenze, Italy; Museo di Storia Naturale, Sezione di Zoologia “La Specola”, Università degli Studi di Firenze. Via Romana 17, 50125, Firenze, Italy; Dipartimento di Biologia, Università degli Studi di Firenze. Via Madonna del Piano 6, 50019, Sesto Fiorentino, Italy; Regione Toscana. Via di Novoli 25, 50127, Firenze, Italy; Universiteit Utrecht. Padualaan 14, 3584 CH Utrecht, Netherlands

## Abstract

Rule violation is critical for biological conservation worldwide. Conventional questionnaires are not suitable to survey these violations and specialized questioning techniques that preserve respondents’ privacy, like the forced-response RRT, have been increasingly adopted by conservationists. However, most of these approaches do not measure multinomial answers and conservationists need a specialized questioning technique for real-world settings where non-compliance could occur in different forms. We developed a multidimensional, statistically-efficient, RRT which is suitable for multinomial answers (mRRT) and which allows researchers to test for respondents’ noncompliance during completion. Then, we applied it to measure the frequency of the various forms of illegal restocking of European catfish from specialized anglers in Italy, developing an operational code for the statistical software R. A total of 75 questionnaires were administered at a large fishing fair in Northern Italy, in winter 2018. Our questionnaires were easily compiled and the multinomial model revealed that around 6% of respondents had moved catfish across public freshwater bodies and private ponds. Future studies should better address their characteristics, and the mRRT could allow for modeling the effect of co-variates over restocking behavior. The multinomial mRRT could be adopted to measure many forms of rule violation in conservation that could take different forms, like various forms of fish restocking or different modes of wildlife persecution.

## 1 Introduction

### 1.1 Noncompliance in conservation

The violation of existing rules governing the commons often leads to social dilemmas and inefficiencies in their utilization (Ostrom, 2009), with devastating effects over the environment, society and human well-being. Famous cases of noncompliance with conservation rules include overfishing (Coll et al., 2008), illegal logging and poaching (Guan et al., 2018; Kideghesho, 2016; van Velden et al., 2018), illegal wildlife trade (Biggs et al., 2017), violation of biosecurity practices (Trinidade Castro et al., 2017) or the use of forbidden chemical products (Perugini et al., 2018). Counteracting non-compliance with existing environmental regulations is therefore a major goal for conservationists worldwide, and many conservation achievements were obtained by setting proper regulations and by better enforcing existing ones, making them effective in practice. Multiple ways are available to manage and foster compliance with environmental regulations. but understanding the actors and drivers of noncompliance is crucial for their choice (Arias, 2015). Questionnaires can be regarded as one of the most flexible and in-depth approaches for this purpose, as they also collect information to better characterize noncompliant subjects. Unfortunately, conventional direct-answer questions are unsuitable to measure noncompliance in conservation, because respondents generally do not answer honestly due to social desirability or the risk of sanctions (Krumpal. 2013; Nuno and St. John, 2015). Despite in principle it is possible to protect respondents by ensuring anonymity through survey administration mode, respondents generally do not trust conventional questionnaires. Even when researchers persuade respondents to answer honestly, privacy protection in conventional questionnaires is essentially up to their goodwill: local authorities could put pressure on them for disclosing information, data could be hacked, or information could be disclosed by research institutions in a top-down fashion. This constitute a danger both for respondents and for conservationists. Researchers who deal with deviant behaviors and sensitive topics in social sciences (e.g. crime, drug addictions), recognized these issues long ago, and nowadays they generally avoid conventional questionnaires.

### 1.2 Specialized questioning techniques

Specialized questioning techniques were developed to overcome the limitations of direct-answer questionnaires, in terms of privacy protection. They include many different methods which protect respondents’ privacy through a series of instructions that make impossible to know the true answer given to a certain question, at the individual level. In a special issue of Biological Conservation, Nuno and St.John (2015) already discussed the various techniques that were available at that time to measure noncompliance in conservation. It is worth noticing that, since 2015, various advances were proposed for the Crosswise Model (Heck et al., 2017; Tu and Hsieh, 2017), for list experiments (Gaia et al., 2018; Krumpal et al., 2018; Perri et al., 2018), for various non-randomized approaches (Liu et al., 2017; Tian et al., 2017), for Respondent Driven Sampling (Heckathorn and Cameron, 2017; Sperandei et al., 2018) and for the Randomized Response Technique (RRT, Cruyff et al., 2016b; John et al., 2018). The RRT is arguably the most adopted specialized questioning technique in various fields of research investigating sensitive behaviors, including conservation biology. The most common of its versions, the forced response design combines a good statistical efficiency and the use of a randomizing device, which make it good for real-life has applications in conservations: thanks to its statistical efficiency researchers could obtain prevalence estimates from a snample of a few hundreds participants and by using a randomizing device they do not need to know the distribution of some nonsensitive characteristics of the target population (Blair et al., 2015). Moreover, the RRT allow for the development of a likelihood function, to test for the effect of covariates over probability to engage into the sensitive behavior (Cruyff et al., 2016a). Traditionally the forced design RRT estimates the prevalence of binomial sensitive questions, but models for ordinal outcomes were developed (Conteh et al., 2015). Recently, Cruyff et al. (2016b) proposed a multidimensional version of the forced design RRT (mRRT) for ordinal outcomes, based on two consecutive randomized questions. The mRRT is more statistically efficient than conventional RRT by about 75%, requiring about 50 respondents to obtain 10% SE about prevalence estimates. Moreover it could account for respondents’ noncompliance with the instructions, correcting prevalence estimates in the light of self-protective answers. However, a mRRT version for multinomial outcomes has never been proposed. This is a considerable limit to its application in conservation biology, where researchers and practitioners are often interested in characterizing not only the magnitude of noncompliance, but also how it occurs. For example a researcher who studies wildlife persecution might be interested in understanding the various ways through which farmers kill problematic wildlife (e.g. poison, trapping, shooting), or an expert of biosecurity might be interested in which at risk behaviors are performed by some target groups. In this paper we will address this gap by implementing a multinomial version of the RRT, altogether with providing the code for its implementation on a statistical software, and by showing its application in the estimation of various forms of illegal fish restocking affecting the spreading of an invasive alien fish in Italy.

**Figure 1:**
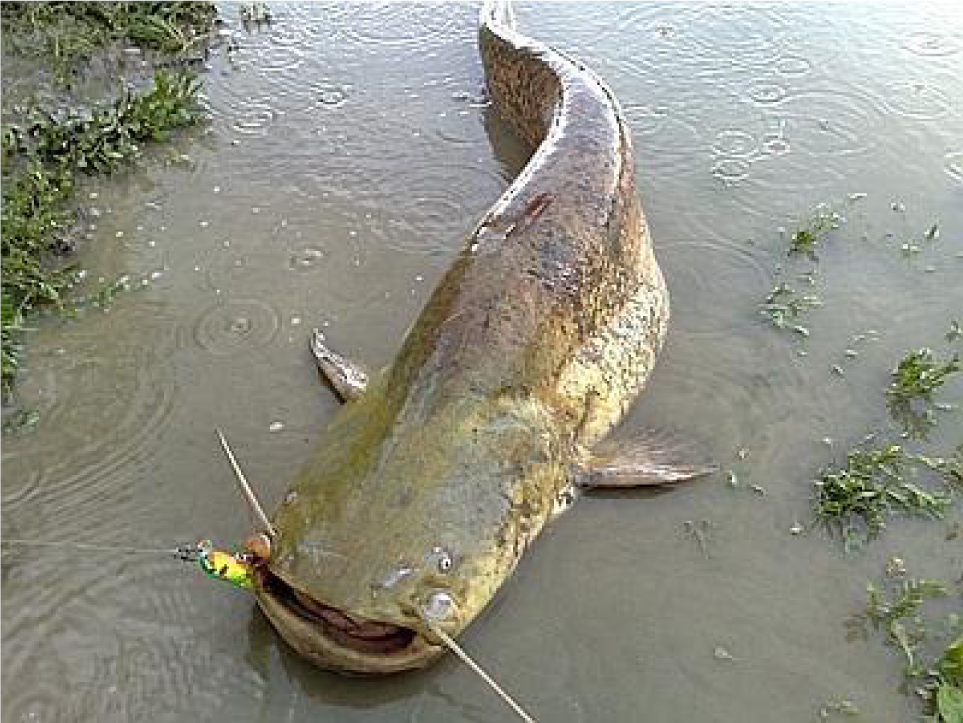
A European catfish from the Arno river (source: Adnkronos)

### 1.3 Case study: the European catfish in Italy

The European catfish (*Silurus glanis*) is an endemic megafish of Eastern Europe and some Central European watersheds. However, it is also a major aquatic invader that established viable populations in France, the UK, Spain and Italy. Despite researchers do not agree about its overall predatory impact, its carnivore diet, coupled with its extremely large size and the high level of environmental degradation reached by some of its introduction ecosystems, raise concerns about the potential consequences of its presence over native freshwater fish communities. The fact that the European catfish is much larger than almost all the other freshwater fish species in its introduction range, makes it a predator for those large species that were not previously predated by any other fish, triggering unpredictable trophic cascades.

The rapid geographical spread of the European catfish outside of its native range in Europe, altogether with its size and its metabolic requirements, indicates that the species is actively restocked by recreational anglers. In facts, despite catfish have a high level of site fidelity and territoriality, they appeared over a very high number of disconnected watersheds across many European countries in a couple of decades. Moreover, their large size stimulated a growing number of recreational anglers to practice trophy fishing in various European countries. The low oxygen requirements of catfish mean that they could be easily stocked in water tubs for a prolonged timespan: this is a crucial prerequisite for moving fished individuals across water bodies, in case anglers wanted to introduce the species to increase their recreational angling experience (Cucherousset et al., 2018). A major gap in our understanding of illegal catfish restocking lies in the fact that multiple forms of restocking exists. While it is known that anglers release catfish across water bodies, it is uncertain whether this activity occurs mostly at public freshwater bodies, at private fishing ponds or in both the environments. This makes catfish restocking a good case study to test specialized questioning techniques for multinomial outcomes, aimed at disentangling the various forms of deviant behavior. In this research we developed a multidimensional Randomized Response Technique to obtain prevalence estimates for a multinomial sensitive behavior, and we applied it to estimate the various forms of illegal catfish restocking over a sample of recreational anglers at a fishing fair in Northern Italy.

## 2 Methods

### 2.1 Questionnaire design and administration

A questionnaire adopting the multidimensional Randomized Response Technique (Cruyff et al., 2016b) was designed. The questionnaire aimed to quantify the frequency of illegal catfish restocking, and its most common forms, among recreational anglers in Italy. In the first page, respondents were asked whether they purchased a fishing permit in 2017, about their age, the type of angling they practiced, the frequency of their catfish angling sessions at private ponds and in public freshwater. Moreover, they were asked to express their beliefs about the impact of European catfish over native fish community and to indicate which management measures they deem to be the best ones for managing catfish populations. Finally, they were asked to indicate the importance of a fishing spot for catfish in their area, for the quality of their angling experience. In the second page of the questionnaire, the multidimensional RRT included two different questions:

- in the **first question** participants were asked: “In the last 12 months, did you restock any freshwater body or fishing pond, with some European catfish you had fished somewhere else?”. Before answering they rolled a 12-faces die. If the outcome was 1 or 2, they were asked to answer ‘Yes’, regardless of what they would have answered instead. Similarly, if the outcome was 11 or 12, they were asked to answer ‘No’. On the other hand, if the outcome was between 3 and 10, respondents were asked to provide an honest answer to the question.
- in the **second question**, respondents were asked “Could you please tell us how many of the following behaviours did you do, in the last 12 months?”. Options included: “I did not restock any catfish, at a fishing spot different from where I fished it”, “I restocked public freshwater bodies with live catfish”, “I restocked one, or more, private fishing ponds with live catfish”, “I restocked both private fishing ponds and public freshwater bodies, with live catfish”. Before answering they rolled again the 12-faces dice: if the outcome was between 1 and 9, they were asked to answer honestly. On the other hand, if the outcome was 10, 11 or 12, respondents were asked to roll again the die and to mark the response option which contained the outcome in a small caption on its side.

The second question quantified the proportion of anglers who engaged in each form of illegal catfish restocking. Available experience indicates that most restocking happens at public freshwater bodies, like streams or rivers. However, we were also interested in exploring how many respondents declared to restock catfish at private ponds, given the economic importance of game fishing. As the two behaviors are not mutually exclusive, we included an option to measure those respondents who restocked both at fishing ponds and at freshwater bodies. The baseline option was designed to include those anglers who did not restocked any catfish in the last 12 month, but to exclude those practicing catch-and-release, a common practice in catfish restocking. A complete English version of the questionnaire is available in Appendix 1.

In February 2018, 75 questionnaires were administered at a sample of anglers at “Pescare show”, the largest fishing fair in Italy. An operator approached anglers in the section about baitcasting and catfish angling, and asked them whether they practiced European catfish angling. Those who answered yes, were asked if they wanted to take part into an anonymous survey about angler behavior. Respondents received the questionnaire and then were left alone during completion. They put the form into a sealed urn, once completed. Given our instructions, respondents rolled the die between 2 and 3 times. Data can be regarded as a mixture of respondents who answered honestly and respondents who were forced to provide a certain answer, on the basis of the outcome on the die (Fig. 2).

**Figure 2:**
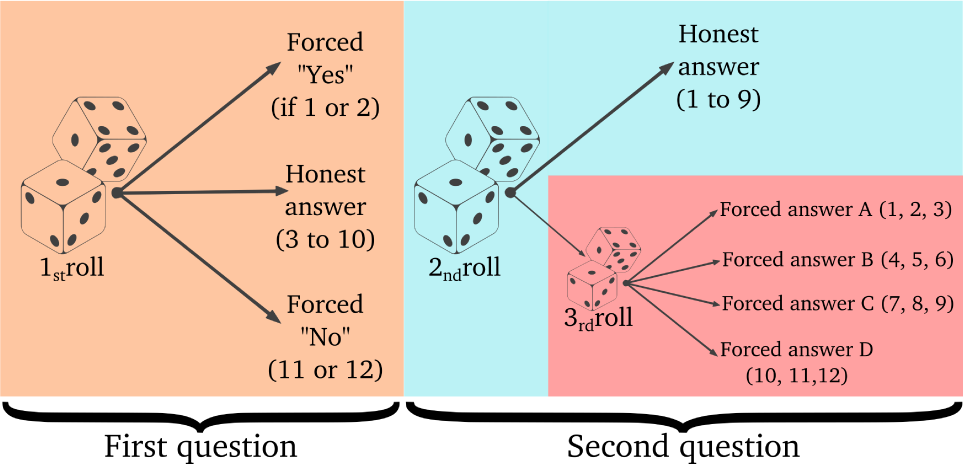
Figure 2: how the mRRT adopted in this study worked in practice.

A complete example of data analysis in R is available in Appendix 2 and a complete dataset is available as a Supplementary Material.

## 3 Results

Questionnaires were piloted over a convenience sample of anglers. Piloting did not reveal any particular issue and questions were generally understood by respondents. The first respondents were told that explanations about the randomized response were available on request. However, participants did not asked for further clarifications, completing the survey in about five minutes and admitting that everything was relatively clear.

Respondents’ age was 40.2 ± 16.7 years (mean ± sd), and 86.7% of them declared to purchase a fishing permit in 2017. Baitcasting was the dominant form of fishing (64%), followed by flyfishing (32%), coarse fishing (0.25%), sea fishing (24%), various unspecified forms of fishing (0.20%) and carpfishing (18.7%). The majority of respondents believed that catfish could have some negative impact over native fish species (64%). 62.7% of respondents believed that catch-and-release was an important measure for catfish management. A good proportion of anglers deemed that having a fishing spot for the European catfish nearby, would not improve their recreational experience (48.8%). Surprisingly, contrary to what they said to recruiters, a good proportion of respondent said that in 2017 they never went fishing European catfish, neither at fishing ponds nor at lakes: we will discuss this point in the following section.

The multinomial randomized response model indicated that the majority of anglers did not restocked any catfish in the last 12 months (91.7%), but also that some respondents restocked catfish at freshwater (6.0%) and that 2.3% of respondents did so at both freshwater and private fishing ponds). No angler seemed to have restocked catfish at private fishing ponds only (Table 1). One advantage of the mRRT is the possibility to model the observations, by assuming that some respondents engage in self-protective strategies, violating the instructions of the RRT. For example respondents could mark the “No” option, regardless of the outcome of the die or their real behavior. This form of self-protection can be obtained by fitting an zeroinflated model and by comparing it with the original model (e.g., through information criteria) observing which one of the two models fits the data better. In this research, we fit a zero-inflated model accounting for respondents noncompliance with the instructions, but we did not notice any improvement over the initial model. We therefore retained the original mRRT model which assumes that respondents complied with the instructions.

**Table 1:** Prevalence estimates and 95% confidence intervals of restocking.

| Behavior | Frequency | 2.5% | 97.5% |
| --- | --- | --- | --- |
| No restocking | 0.917 | 0.817 | 0.990 |
| Freshwater | 0.060 | 0.000 | 0.146 |
| Private ponds | 0.000 | 0.000 | 0.000 |
| Freshwater and ponds | 0.023 | 0.000 | 0.086 |

## 4 Discussion, conclusions and research perspectives

Our research is the first where a multinomial version of the multidimensional RRT was adopted. Our findings are controversial and they must considered absolutely preliminary. In facts, while practicing catfish angling was a prerequisite for being recruited into our study, most respondents told that they did not fish any European catfish in the 2017. Moreover, the presence of a nearby fishing spot with European catfish was not perceived to be crucial for the quality of recreational angling. These two points could indicate that our respondents could have been recreational anglers who were not specialized gamefishers. Alternatively they might also indicate that respondents defined themselves as catfish anglers but that they looked suspiciously to researchers and did not disclose any information about themselves, when approached. Finally, respondents could have been tempted to present themselves as regular catfish anglers, to be rewarded for their answers, a common practice in survey research, especially in marketing studies conducted at public events (Singer and Ye, 2013). We believe that future studies should focus on specialized catfish anglers by means of online surveys promoted on social networks and fishing websites. Despite developing online versions of the RRT is not straightforward, online administration could be very useful for intercepting highly-specialized recreational anglers who invest time and money on dedicated fishing equipment for catfish angling. Under a practical viewpoints, our findings indicate that a proportion of recreational anglers in Italy, seems to illegally restock catfish. However, we are cautious about them and we do not generalize, for the various reasons we mentioned above. Future studies should focus on underlining the most common forms of catfish restocking, to better understand both the sites of introduction and the potential vectors. This would be useful for counteracting the spread of European catfish in Italian freshwater. For example, determining where catfish are introduced by anglers, could enable conservationists to optimize monitoring efforts, selecting those areas that have to be monitored. Establishing that catfish are not released at private fishing ponds means that these places could not be monitored, as they are not hotspots of catfish introductions.

Our findings are way more interesting in a methodological perspective: we believe that the multinomial multidimensional RRT could be very useful for conservationists worldwide. In our case study, we used it to estimate the prevalence of various pathways of biological invasions. But, its potential use in the management of natural resources and biodiversity conservation could extend to a wide range of behaviours, like wildlife persecution. In a previous work Cerri et al. (2017) estimated the frequency of illegal wildlife killing among recreational farmers, in Central Italy, distinguishing between four groups of wildlife by means of four different RRT questions. The mRRT could have been very useful to combine these questions into a single one. Moreover, the mRRT could have been adopted to design a questionnaire aimed at distinguishing the various methods adopted to kill wildlife. This latter option could be particularly useful to measure the techniques adopted in carnivore persecution (St. John et al., 2011, 2015), which often implies the use of poison (Santangeli et al., 2016), but also traps or shooting. Implementing a multinomial RRT with the multidimensional design offers two other advantages, compared to conventional RRT. Firstly, it is statistically efficient: while the conventional forced-response design for binomial outcomes requires more than 800 observations, to obtain a 95% confidence interval, mRRT requires much smaller samples. Cruyff et al. (2016b), for their case study, showed that sample size was reduced by 75%, a remarkable decrease that is certainly compatible with most field studies in conservation. Of course, the statistical power of the technique depends on the probability of a truthful response and on the number of answers to the second question. The lower the number of response options, the higher the statistical efficiency, as the multinomial model needs to estimate a smaller number of parameters. The second advantage lies in the possibility to test for respondents noncompliance. This opportunity is important both to obtain accurate prevalence estimates, and to measure data quality in general. In our model, we included the possibility that respondents violated the instructions by selecting the safe answer option. However, various forms of noncompliance exist and they should be incorporated in the model on the basis of data at hand and substantive theory. Further insights on how to implement models accounting for noncompliance could be found in Cruyff et al. (2016b). Finally, it is worth noticing that John et al. (2017) conducted various experiments to test in which conditions the RRT outperforms or underperforms conventional questionnaires. Their results indicate that respondents care about the misinterpretation of their answers and that some particular response formats should be adopted. We believe that future experimental studies should be replicated for the mRRT, which is probably more complex and cognitively demanding than traditional RRT designs, to test how it affects respondents during the answering process.

## Appendix 1 Complete questionnaire

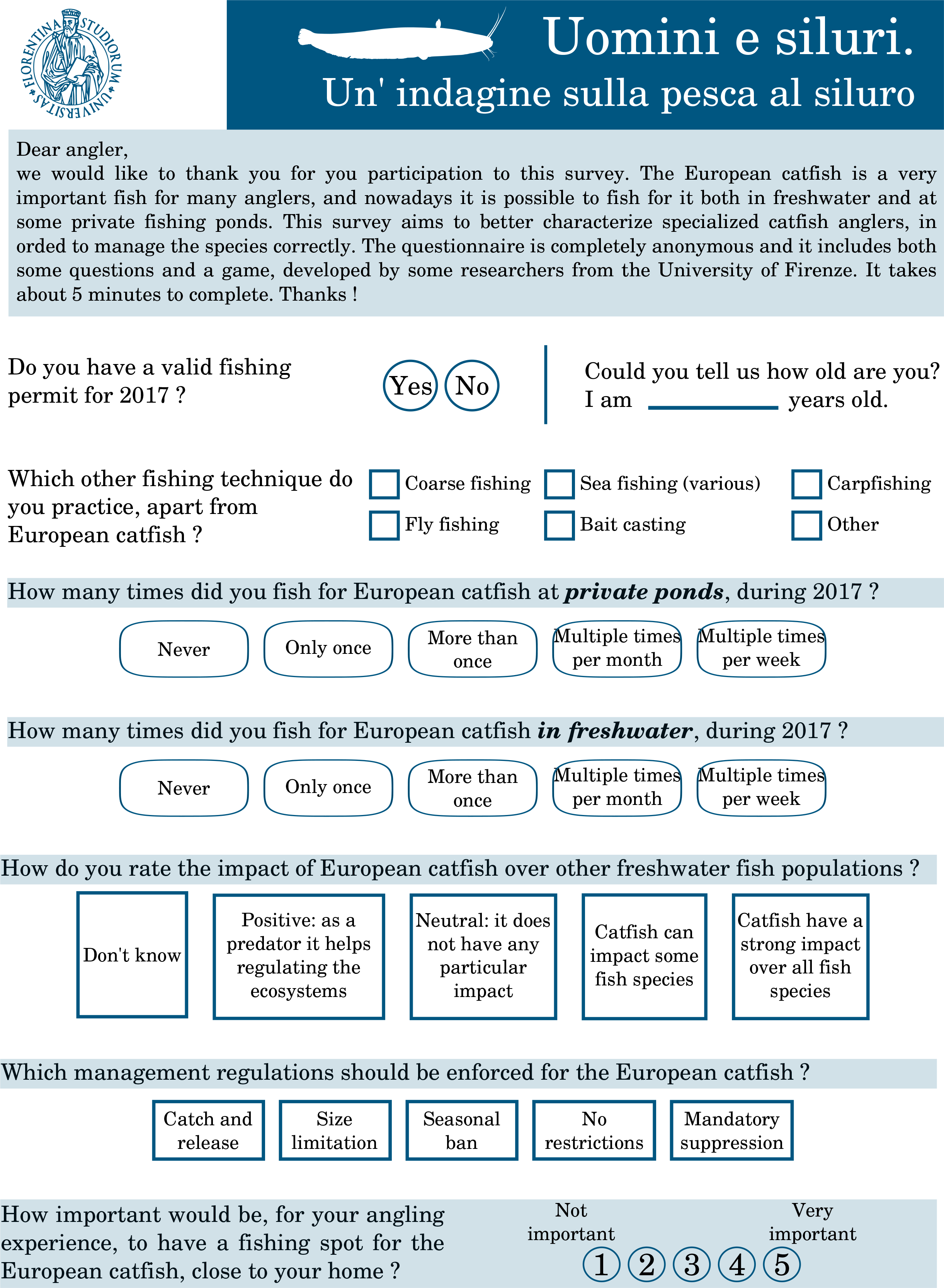

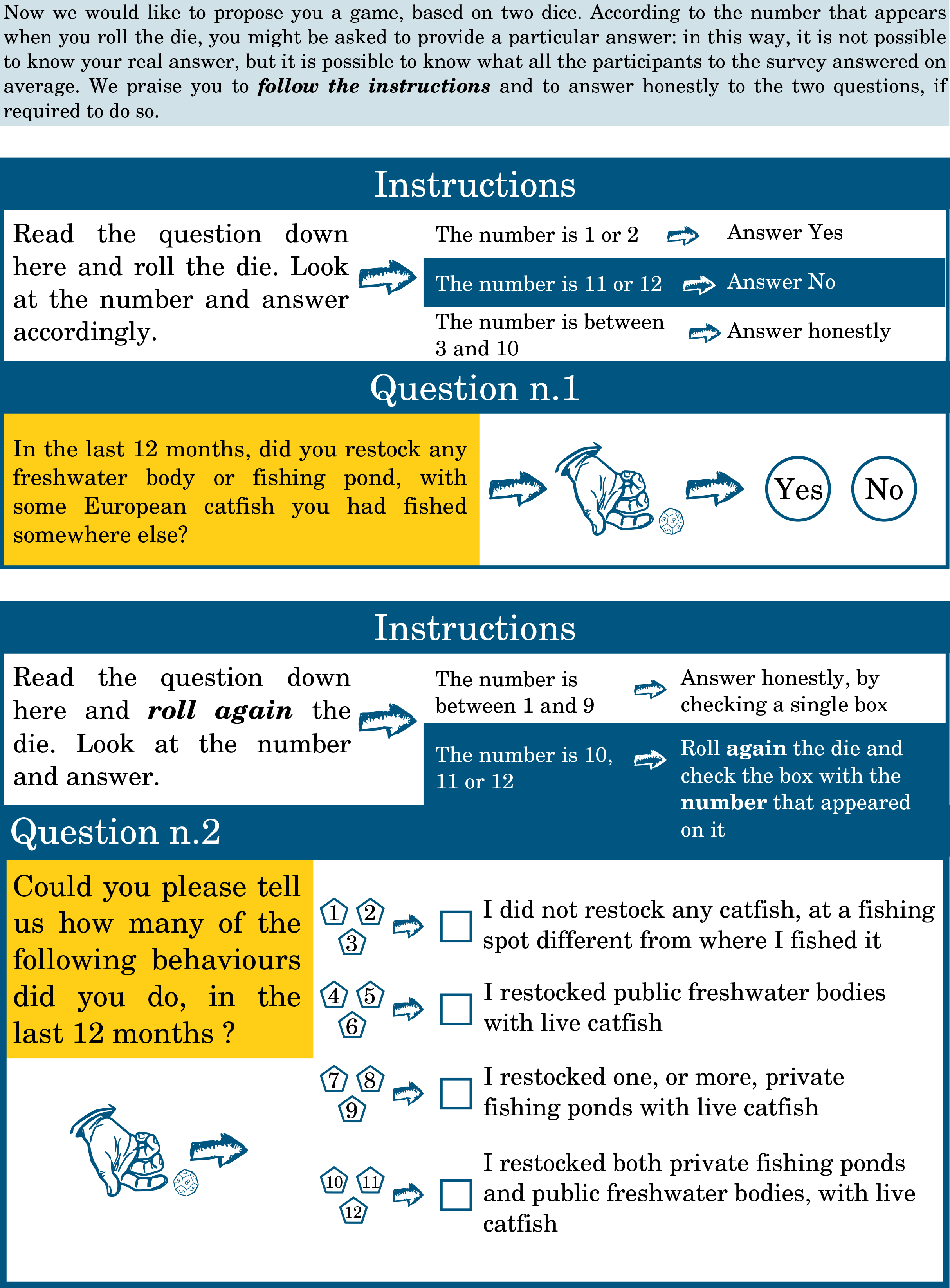

## Appendix 2 Reproducible R code

The following code is entirely reproducible with the statistical software R. It does not require any particular package, apart those which are automatically included in the software. Let’s import the dataset and label the combinations of the first and second randomized questions.

~~~
nobs <− c(36,10,7,6,11,3,0,2) *# observed data*
names(nobs) <− c(“0A”, “0B”, “0C”, “0D”, “1A”, “1B”, “1C”, “1D”) *#Naming combinations*
n <− sum(nobs)
~~~

### Misclassification matrix

The misclassification matrix is the basis of the Randomized Response model. The conditional misclassification probabilities are computed as the products the conditional misclassification probabilities for the separate questions. For example, if the dominant restocking behavior is C, then the probability to answer yes to the first question is 5/6, which is the probability that the outcome of the die is 1 to 10 (resulting in either a forced or an honest “Yes”). The probability of answering C to the second question is equal to 13/16, which is the sum of the probability 3/4 of an honest “Yes” answer to the first question, and 1/16 which is the the product of the probabilities of throwing 10, 11 or 12 with the first die (1/4), and having to answer answering C on the second die (1/4). Hence the conditional misclassification probability P (yes, C|C) = 5/6 × 13/16 = 0.6771.

~~~
dQ1 <− 2/3 *# probability of truthful response on answer Q1 and Q2*
dQ2 <− 3/4 *# probability of truthful response on answer Q1 and Q2*
b <− 1/6 *# probability of forced“Yes” to Q1*
c <− 1/6 *# probability of forced“No” to Q1*
e <− 1/16 *# probability of forced A, B, C, D*
~~~

~~~
P1 <− matrix(c(dQ1+c,b,c,dQ1+b),2,2) *#matrix(c(ptrue+pno,pyes,pno,ptrue+pyes),2,2)*
P2 <− matrix(e,4,4) *#matrix(e,4,4)*
diag(P2) <− dQ2+e
P <− cbind(P1[,1]%x%P2[,1],P1[,2]%x%P2[,2:4])
~~~

### Writing down the model and compute the 95% confidence interval

An approxinmate confidence interval can be obtained with a nonparametric bootstrap. The procedure is to draw with replacement 1, 000 random samples of size n = 75 from the observed responses, save the prevalence estimates and compute the 2.5 and 97.5 percentiles for the prevalence of each restocking behavior.

~~~
lg <− function(v)exp(c(0,v))/(1+sum(exp(v)))
RR <− function(par, data){−t(data)%*%log(P%*%lg(par))}
est <− optim(rep(−3, 3), RR, hessian=T, method=“BFGS”, data=nobs)
phat <− matrix(lg(est$par), 4, dimnames=list(0:3, “phat”))
~~~

~~~
boot.phat <− matrix(0, 1000, 4)
set.seed(100)
for(i in 1:1000){
  boot.sample <− sample(1:8, size=sum(nobs), replace=T, prob=nobs/sum(nobs))
  boot.obs <− data.frame(table(factor(boot.sample, levels=1:8)))$Freq
  boot.est <− optim(rep(−3, 3), RR, hessian=T, method=“BFGS”, data=boot.obs)
  boot.phat[i,] <− lg(boot.est$par)
}
~~~

We can therefore estimate coefficients and their 95% confidence interval.

~~~
round(cbind(phat, t(apply(boot.phat, 2, function(x)quantile(x, probs=c(0.025, .975))))), 3)
~~~

~~~
##    phat  2.5% 97.5%
## 0 0.917 0.817 0.990
## 1 0.060 0.000 0.146
## 2 0.000 0.000 0.000
## 3 0.023 0.000 0.086
~~~

